# Amelioration of L-DOPA-induced dyskinesia with vitamin D_3_ in Parkinsonian mice model

**DOI:** 10.1101/2021.09.13.459937

**Authors:** Adedamola Aminat Bayo-Olugbami, Abdulrazaq Bidemi Nafiu, Abdulbasit Amin, Olalekan Michael Ogundele, Charles C. Lee, Bamidele Victor Owoyele

## Abstract

L-DOPA Induced Dyskinesia (LID) is associated with prolonged L-DOPA therapy. Vitamin-D receptor modulation improves motor-cognitive deficit in experimental LID Parkinsonism. Therefore, in this study, we investigated the mechanism underlying the anti-dyskinetic potential of Vitamin D3 (VD_3_). Dyskinesia was induced by chronic L-DOPA administration in 6-OHDA lesioned male C57BL6 mice. The experimental groups (Dyskinesia, Dyskinesia/VD_3_, and Dyskinesia/Amantadine) and controls were challenged with L-DOPA to determine the abnormal involuntary movements (AIMs) score during 14 days of VD_3_ (30 mg/kg) or Amantadine (40 mg/kg) treatment. Global behavioral Axial, Limb & Orolingual (ALO) AIMs were scored for 1 min at every 20 mins interval, over a duration of 100 mins on days 1,3,7,11 and 14 of treatment. Thereafter, brain samples were collected and processed for immunoblotting to assess striatal expression of tyrosine hydroxylase (TH), monoamine oxidase (MAO), cathecol-o-methyl transferase (COMT), dopamine decarboxylase (DDC), CD11b, BAX, P47phox, and IL-1β. VD_3_ significantly attenuated ALO AIMs only on days 11 & 14, with maximal reduction of 32.7% compared with dyskinetic mice but had no effect on days 1, 3 & 7, while amantadine decreased AIMs all through days 1 to 14 with maximal reduction of 64.5%. TH and MAO-B expression were not significantly different across the groups. DDC was significantly suppressed in dyskinetic mice *vs* control (p<0.001) but remained unchanged in VD_3_ mice *vs* dyskinetic mice. COMT was upregulated in the dyskinetic group *vs* control (p<0.01) and attenuated in VD_3_ mice (p<0.05) compared to the dyskinetic group. Interestingly, VD_3_ inhibited significantly (p<0.01) oxidative stress (p47phox), apoptosis (BAX), inflammation (IL-1β), and microglial activation (CD11b) in dyskinetic mice. Overall, we find that the anti-dyskinetic effects of VD_3_ is associated with modulation of striatal oxidative stress, microglial responses, inflammation, and apoptotic signaling.

**Impact statement:** There are evidences showing that VD_3_ supplementation improves motor disorders, including Parkinson’s disease. We hypothesized that VD_3_ could improve LID, an abnormal involuntary movement that results from prolonged L-DOPA therapy in the management of PD. We have demonstrated the novel anti-dyskinetic effect of VD_3_ and associated mechanistic factors in a mouse model of L-DOPA Induced Dyskinesia (LID), which identifies promising targets for dyskinesia therapy.

## Introduction

Development of treatment-induced motor complications constitutes a major problem in the long-term management of Parkinson’s disease (PD). Particularly, L-DOPA-induced dyskinesia (LID) poses a significant challenge. L-DOPA, the predominant drug for alleviating the motor deficit associated with PD, is fraught with uncontrolled abnormal movements termed LIDs.^1, 2^ In addition, prolonged use of L-DOPA by PD patients causes increase in dopamine (DA) toxicity,^3^ oxidative stress^4-6^ and induction of inflammatory responses.^7^ Thus, monitoring inflammatory factors and apoptotic proteins has been recommended in L-DOPA therapy.^8^ Most PD patients develop dyskinesia in less than a decade from onset of treatment.^9, 10^ Despite the clinical importance of this side effect, the mechanisms underlying the generation of LIDs are not fully understood. On the one hand, the degree of dopamine loss and the dose of L-DOPA, rather than the duration of L-DOPA treatment, may correlate with the development and severity of dyskinesia.^11, 12^ Conversely, dyskinesia may also result from prolonged use of L-DOPA, which causes the release of dopamine in the brain without a corresponding uptake and break down, in effect over stimulating D1 receptors leading to dyskinesia.^13^

Animal models in which neurotoxins were used to cause unilateral selective loss of substantia nigra neurons, typical of PD, have been used to assess LIDs ^14-16^ and the efficacy of standard and novel therapeutic treatments. These models suggest that dyskinesias are associated with enhanced G protein-mediated signaling at dopamine receptors,^17-19^potentially leading to changes in gene expression and uncontrolled neuronal excitability.^17-22^Therefore, therapeutic strategies that can moderate calcium signaling and neuronal excitability while maintaining normal movement may be an ideal way to eliminate DA receptor-associated dyskinesias. To moderate this uncontrolled signaling or neuronal excitability, several approaches have been explored such as reducing D1 receptors surface expressions,^23, 24^ dampening overactive intracellular signaling^25, 26^ and inhibiting α2A,^27^ mGluR5 ^28, 29^ or NMDA receptors. ^23, 30^ Although these targets have clinical potential, several drugs designed for these targets have either failed clinical trials or have the potential to affect other key CNS physiological processes.^31^

In this regard, the vitamin D receptor (VDR) appears crucial for brain development and function in health and disease, potentially mediating up-regulation of glutamate receptors, calcium-induced excitotoxicity and reactive oxygen species (ROS) (all of which are Ca^2+^ dependent toxicities and central to the cause and progression of PD and dyskinesia).^32^ Therefore, targeting a calcium-controlling receptor, VDR with a large presence in the striatum, can be an effective approach to treating dyskinesia.

Vitamin D (VD) is a steroid that is capable of up-regulating genes involved in protein synthesis and has been found to improve survival in cells by facilitating repair. The importance of VDR, a major Ca^2+^ controlling receptor, with a large presence in the striatum is still underexplored. Although several reports have implicated VD in some neurological disorders, its effect on L-DOPA-induced dyskinesia has not been explored. Other studies have favored experiments on the effect of Vitamin E in reducing the threshold of Tardive dyskinesia (TD) (a side effect of prolonged use of antipsychotic drugs). The anti-dyskinetic effect of Vitamin E, though in TD was attributed to its antioxidant properties and its role in radical detoxification. ^33, 34^ Additionally, oxidative stress, microglia activation, neuro-inflammation and apoptotic signaling have been implicated in the pathophysiology of dyskinesia.^35, 36^ Since VD is reportedly involved in the regulation of these processes both in disease and non-pathological state, investigating its role may hold a promising therapeutic effects in the treatment of dyskinesia. Similarly, VD_3_ deficiency has been linked to the cause and progression of various movement disorders,^37^ including PD, wherein it was reported to positively modulate and improve neurotransmission as well as behavioral deficits in a mouse model of PD.^38^ By virtue of its roles in radical detoxification, Ca^2+^-related signaling and general brain health, VD_3_ is a potential therapeutic agent in attenuating dyskinesia. Currently, no study has investigated the impact of VD_3_ on proteins involved in dopamine metabolism, behavioral alteration and oxidative stress in animal model of LID. Therefore, we assessed how behavioral (using abnormal involuntary movements (AIMs) rating scale), biochemical and molecular (using immunoblotting) responses are altered in mouse models of dyskinesia induced by chronic L-DOPA administration in *unilateral* 6-hyroxydopamine (6-OHDA) lesioned mice and the impact of VD_3_ on these alterations was evaluated.

## Materials and methods

### Drugs

All drugs used were of analytical standard. 6-OHDA and Pargyline hydrochloride were purchased from Sigma Aldrich (Sigma Chemical, St. Louis, MO). L-DOPA, Amantadine and Benserazide were obtained from TCI, America (North Harborgate St., Portland, US) while Apomorphine, Desipramine hydrochloride and Cholecalciferol (VD_3_) were products of USPharmacopeia (Rockville, Maryland, US). Drugs were freshly prepared in saline or saline containing 0.02% ascorbic acid and used within 3 hours of preparation.

### Experimental animals

Age-matched (15 weeks), adult male C57BL6 mice with an average weight of 30.3 g were procured from the Jackson laboratory (Bar Harbour, ME). All studies were performed according to NIH guidelines for animal care and use and as approved by the Animal Care and Use Committee of School of Veterinary Medicine, Louisiana State University with protocol number, 16-015 and University of Ilorin Ethical Review Committee with approval number, UERC/ASN/2017/738. Mice were housed at a maximum of five per cage, under 12 h light/ dark cycle and given food and water *ad libitum*.

### Experimental schedule/animal grouping

Thirty mice were lesioned by right *intrastriatal* injection of 6-OHDA. Two weeks later, twenty-five 6-OHDA lesioned mice were successfully screened and selected using a battery of behavioral tests. An additional group of mice (n=6) was sham lesioned (control). Dyskinesia in the 6-OHDA lesioned mice was induced by chronic L-DOPA administration (20 mg/kg combined with 12.5 mg/kg benserazide) once daily for 10 days to induce gradual development and stable degree of AIMs. During the development of dyskinesia, AIMs were monitored and scored for 1 min at every 20 mins interval, over a duration of 120 mins following L-DOPA challenge on days 1, 3, 6, 8 and 10. ^14, 16, 39^ Each AIMs subtypes: Axial, Limb, Orolingual was rated on frequency and amplitude (Lundblad *et al*., 2007). Eighteen dyskinetic mice with global ALO AIMs (Sum of the products of frequency and amplitude of ALO AIMs) of ≥100 were randomly sorted into 3 groups: Dyskinesia, Dyskinesia/VD, Dyskinesia/Amantadine (n = 6) and a sham-lesioned (control) group (n=6) was added. VD_3_ (30 mg/kg, *subcutaneously*; *s*.*c*.) or amantadine (40 mg/kg, *intraperitoneally*; *i*.*p*.) was administered for 14 days. To quantify the effect of VD_3_ or amantadine on dyskinesia, each group was challenged with L-DOPA. Each ALO AIMs was scored for 1 min at every 20 mins interval, over a duration of 100 mins after L-DOPA injection on days 1,3,7,11 and 14 of VD_3_ or amantadine administration. After behavioral characterization, the mice were euthanized, brains collected and processed for *striatal* expression of tyrosine hydroxylase (TH), monoamine oxidase-B (MAO-B), cathecol-o-methyltransferase (COMT), dopamine decarboxylase (DDC), CD11b, BAX, P47phox, and IL-1β in the *striatum* using western blot assay (Figure 1).

**Figure 1:**
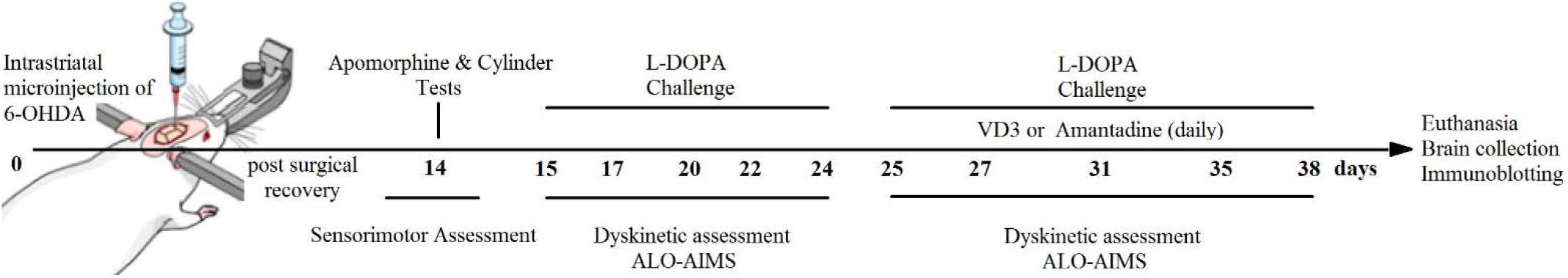
Experimental design showing the treatment time line in L-DOPA-induced dyskinesia following VD_3_ administration. ALO AIMs; Axial-Limb-Orolingual Abnormal Involuntary Movements.

### Unilateral 6-OHDA lesioning

Experimental Parkinsonism was achieved as previously described ^14^ through *unilateral* injection of (6-OHDA, 3µg/2µL) stereotaxically into the *striatum*. Mice were anaesthetized with ketamine/xylazine (80/5mg/kg *i*.*p*.). Saline (0.9%) containing 0.02% ascorbic acid was used to dissolve 6-OHDA and injected in the *striatum* according to the following co-ordinates from bregma: *anteroposterior* 1.1mm, *mediolateral* 1.5 mm, and *dorsoventral* 3.0 mm below *dura*. To increase the selectivity and efficacy of 6-OHDA lesion, mice were pretreated with desipramine (28 mg/kg, *i*.*p*.) and pargyline (6.15 mg/kg, *i*.*p*.) to reduce 6-OHDA-induced noradrenaline/serotonin depletion and enhance the sensitivity of dopaminergic terminals to 6-OHDA 30 mins prior to surgery. They were closely monitored for recovery after which mice were subjected to thorough post-surgical care to minimize mortality.

### Animal screening and behavioral studies

Sensorimotor assessment of the *unilateral* 6-hydroxydopamine-lesioned mice was carried out using drug-based (apomorphine) and a non-drug based (cylinder) ^40^ behavioural tests for the evaluation of degree of lesion. The assessment was performed 14 days after 6-OHDA lesion when mice had achieved a complete post-surgical recovery, ^41^ with a stable and maximal level of dopamine depletion. ^42^ The selected 6-OHDA lesion mice have ≥ 50 contralateral turns/30 mins in apomorphine test and less than 45% contralateral wall contact in cylinder test. ^40^

### Apomorphine-induced rotational response

To determine the efficiency of 6-OHDA-induced *striatal* lesion and dopamine reduction. Rotational response of each mouse to apomorphine (1 mg/kg *i*.*p*.) was observed two weeks after lesion. Each mouse was placed in a cylinder of 30 cm diameter, placed 45 cm below the recording camera. After 3 mins habituation period, rotational movements were recorded over 30 mins time frame. Both contra- and ipsilateral full body rotation relative to the lesioned side were counted and compared with the sham control, i.e. greater contralateral rotation indicative of greater parkinsonism ^42^. The proportion of contralateral rotation was calculated as a percentage of net (contralateral and ipsilateral) rotational behavior.

### Cylinder test

The cylinder test was used to assess spontaneous independent forelimb lateralization, taking advantage of the natural exploratory instinct of rodents to a novel environment by using the forelimb to support the body against the walls of a cylindrical enclosure. Mice were placed individually in a glass cylinder (12 cm diameter, 14 cm height) and video recorded for 10 mins. Mice were not allowed to become habituated to the cylinder prior to the test. The number of wall touches (contacts with fully extended digits) executed independently with the ipsilateral and the contralateral forepaw were counted. Simultaneous paw touches were excluded from the analysis. Data was expressed as a percentage of contralateral paw touches calculated as: (contralateral) / (ipsilateral + contralateral) paw touches × 100.

### L-DOPA-induced dyskinesia and abnormal involuntary movement ratings

Dyskinesia was induced in the animals by chronic administration of L-DOPA (20 mg/kg; *i*.*p*.) and benserazide (12.5 mg/kg; *s*.*c*.) once daily for 10 days. ^18, 31^ Quantification of LIDs by abnormal involuntary movements (AIMs) was carried out as previously described. ^14, 39, 43^ Briefly, mice were observed individually for 1 min every 20 mins during 1-2 h that followed L-DOPA injection. Dyskinetic movements were classified based on their topographic distribution into three subtypes: (i) axial AIM, that is twisted posture or choreiform twisting of the neck and upper body towards the side contralateral to the lesion; (ii) forelimb AIM, that is, jerky or dystonic movements of the contralateral paw; (iii) orolingual AIM, that is, orofacial muscle twitching, empty masticatory movements and contralateral tongue protrusion. AIMs subtypes were scored using a validated rating scale by a blinded trained investigator. Each AIM subtype was rated on frequency and amplitude scales from 0 to 4 as described by Cenci and Lundblad. ^43^ Axial, limb and orolingual (ALO) AIMs were presented together as global AIMs score and also as separated items per session (sum of the products of amplitude and frequency scores from all monitoring periods). ^44^

### Western blotting and protein quantification

Western blot analyses were performed on the mice brain tissue homogenates on the last day of the treatment regimen. Briefly, striatum was rapidly dissected from mouse brains on ice and tissue samples were immediately lysed in ice-cold RIPA buffer solution. Brain tissue lysate (20 µl) containing 20 µg of protein was processed for SDS-PAGE electrophoresis. After subsequent western blotting (wet transfer), polyvinylidene fluoride membrane (PVDF) was incubated in tris-buffered saline with 0.01% Tween 20 (TBST) for 15 mins at room temperature. Subsequently, the membrane was blocked in 3% bovine serum albumin (prepared in TBST) for 50 min at room temperature. The protein of interest, and control (GAPDH) were detected using the following primary antibodies; anti-TH (Cell Signaling-#2792S), anti-COMT (Cell Signaling- #9102S), anti-MAO-B (Abcam-ab175136), anti-DDC (Cell signaling-#13561S), anti-GAPDH (Cell Signaling-#2118S), anti-BAX (Cell signaling-# 14796S), anti-CD11b (Abcam-ab75476), anti-IL-1β (Cell signaling-#2837S) and anti-p47phox (Enzo life Sci-#07071127A). All primary antibodies were diluted in the blocking solution at 1:500-1,000. Subsequently, the primary antibodies were detected using HRP-conjugated secondary antibodies (goat anti-rabbit#65-6120 and goat anti-mouse-#65-6520; Invitrogen; dilution of 1: 5,000-10,000) following which the blots were developed using enhanced chemiluminescence substrate (Thermo Fisher-#34579).

### Statistical analysis

AIMs rating was expressed as ALO score (magnitude x amplitude). Results were reported as means ± SEM and analyzed using Graph Pad Prism (Version 6.0). Differences between groups were determined by one-way ANOVA followed by Tukey’s test for *post-hoc* comparisons. The level of significance was considered at *p*< 0.05.

## Results

### Development and progression of AIMs following chronic L-DOPA administration

*Unilateral* 6-OHDA lesioned mice, chronically treated with L-DOPA (20 mg/kg plus 12.5 mg/kg of benserazide), developed a stable degree of dyskinesia at the sixth day of treatment, scoring the maximal cumulative ALO AIMs value at the 8th day (Figure 2A). Considering the development of individual ALO AIMs, the appearance of axial, limb and orolingual behavioral AIMs was gradual and progressive having a similar temporal profile and reaching a similar level of intensity over the 10-day treatment. Orolingual AIM subtype had the least score while axial appearance was the most represented with the highest score (Figure 2B). L-DOPA caused the appearance of dyskinetic movement already at 20 mins after injection. The intensity of dyskinesia remained stable at maximal levels up to 60 mins after injection, after which AIMs tended to decline (Figure 2C).

**Figure 2:**
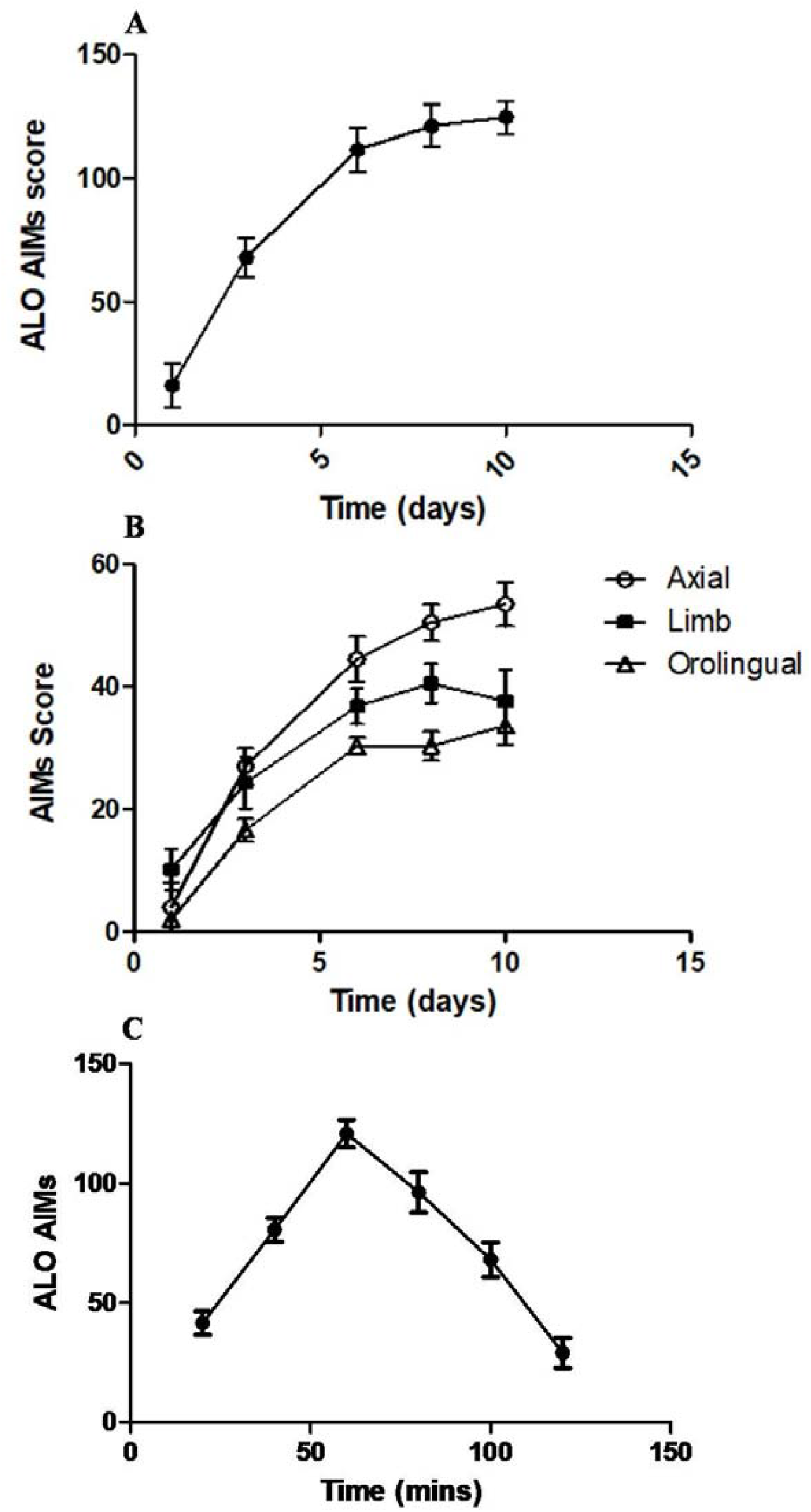
Development and progression of AIMs following chronic L-DOPA administration in 6-OHDA lesioned mice. Cumulative ALO AIMs in 10 days (A); development of individual ALO AIMs, the appearance of axial, limb and orolingual behavioural AIMs (B) and Cumulative ALO AIMs in 2 hours (C).

### VD_3_ attenuated L-DOPA-induced AIMs in dyskinetic mice

Control mice showed no response to any of the ALO AIMs subtype. In the L-DOPA-induced dyskinesia model, chronic administration of L-DOPA caused a significant development of dyskinetic movement assessed by the appearance of ALO AIMs and represented as the global ALO AIM score (Figure 3). To determine the impact of VD_3_ on LID, dyskinetic mice were treated with 30 mg/kg of VD_3_ consecutively for 14 days with the behavioral AIMs response assessed at days 1,3,7,11 & 14. The efficacy of VD_3_ was also compared with amantadine, a common clinical anti-dyskinetic agent. A dose of 40 mg/kg was used because it was reportedly effective in attenuating ALO AIMs in rats and mice without affecting the locomotive component of AIMs. ^39, 45^ This is considered as a marker of the therapeutic effect of L-DOPA. ^46^

**Figure 3:**
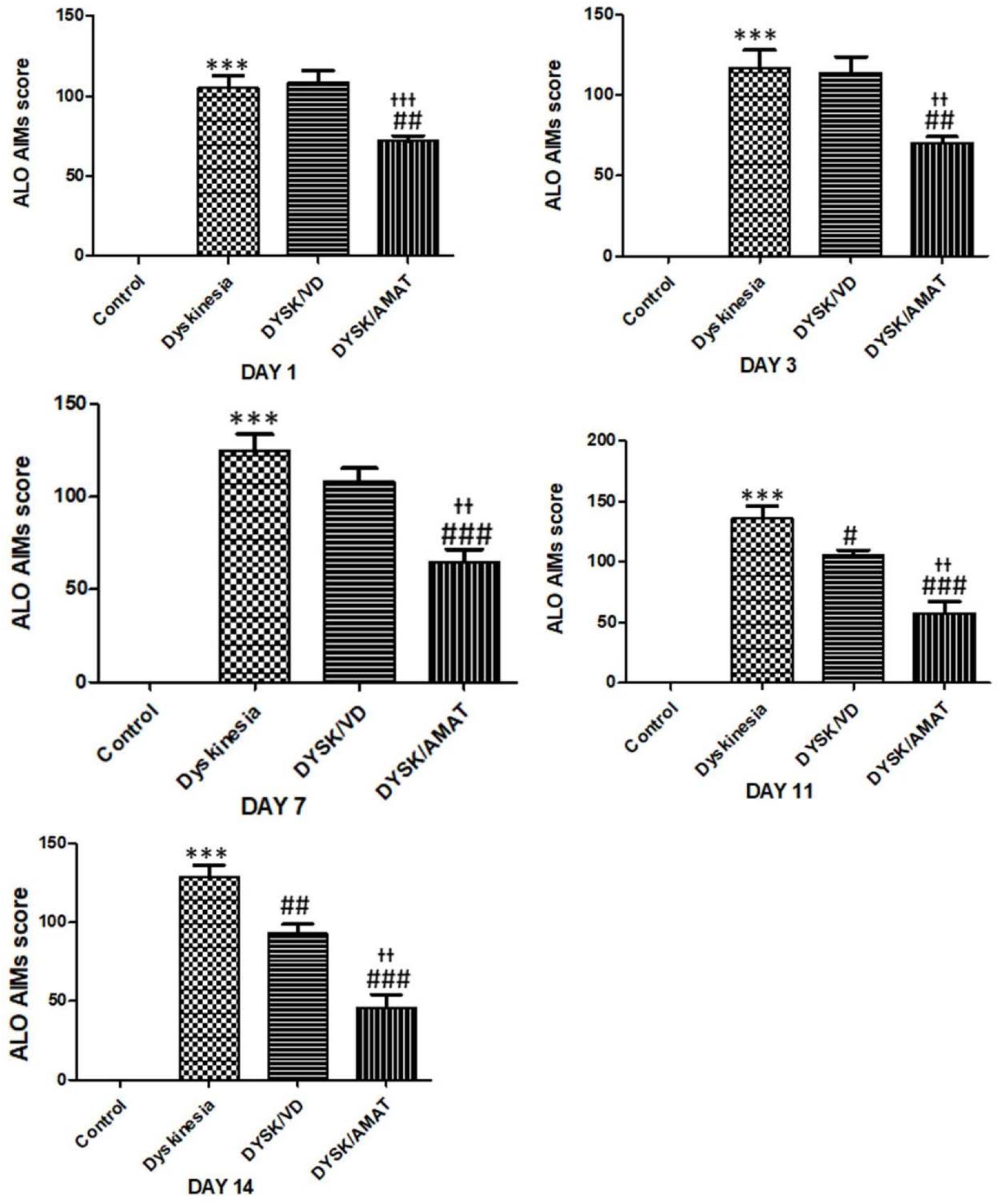
VD_3_ attenuated L-DOPA-induced AIMs in dyskinetic mice. Treatment with VD_3_ did not significantly alter ALO AIMs behaviour in the first seven days (day 1, 3 and 7); improved dyskinesia by reducing the abnormal involuntary behaviours (on day 11 and 14). DYSK/VD; dyskinetic mice that received VD_3_, DYSK/AMAT; dyskinetic mice treated with amantadine. ^***^p<0.001 *vs* control; ^#^p<0.05, ^##^p<0.01, ^###^p<0.001 *vs* DYSK, ^++^p<0.01, ^+++^p<0.01 vs DYSK/VD.

As shown in Figure 3, administration of VD_3_ (30 min before L-DOPA challenge on each day of AIMs assessment) caused no significant reduction in dyskinetic movements at days 1, 3, & 7 with percentage AIMs reduction of 0%, 2.2% & 13.2% respectively. Conversely, it produced a significant reduction in global AIMs on days 11 & 14, which corresponds to 22% and 32.7% reduction in dyskinetic movements respectively. Intervention with amantadine progressively attenuated dyskinesia as depicted by a reduction in the percentage ALO AIMs: day 1 (31.4%,), day 3 (39.5%), day 7 (48%), day 11 (57.5%) and day 14 (64.5%) when compared with dyskinetic mice (Figure 3). Comparing the anti-dyskinetic efficacy of VD_3_ with amantadine showed that amantadine significantly attenuated dyskinetic movements from day 1 to 14 while the anti-dyskinetic effect of VD_3_ was observed between day 11 and 14. As such, it appears as though theVD_3_ associated decline in L-DOPA-induced abnormal involuntary movements was lesser than amantadine (Figure 3). Day 1: F (3, 23) =81.48, p<0.0001; Day 3: F (3, 23) =48.94, p<0.0001; Day 7: F (3, 23) = 67.49, p<0.0001; Day 11: F (3, 23) = 62.39, p<0.0001; Day 14: F (3,23) =77.28, p<0.0001.

### Dyskinetic mice show no alteration in enzymatic regulation of dopamine metabolism

Serotonergic signaling has been implicated as the main underlying cause of uncontrolled DA release in dyskinesia and consequently, the abnormal involuntary movements. In order to examine the potential involvement of the conventional pathway of DA metabolism, the expression of DA synthetic (TH and DDC) and catabolic (MAO-B and COMT) enzymes in the striatum was assessed using western blot analysis.

Expression of TH, the rate limiting enzyme in the synthesis of dopamine, was not significantly different in dyskinetic mice compared with control group (Figure 4A: p=0.48). A similar result was observed for MAO-B (Figure 4E). An increase in the activity of DDC positively affects DA release, but on the contrary, our result showed that DDC expression was significantly lower in the dyskinesia group compared to control mice (Figure 4B: p=0.0002). Furthermore, the activity of COMT was significantly increased in dyskinesia mice compared with control (Figure 4F: p=0.006). As such, if the level of DA in dyskinesia were to be from the conventional pathway of DA metabolism, then COMT level should have been reduced or unaltered that is, DA break down ought to be inhibited thereby enhancing DA availability. In contrast, the reverse was observed which did not support DA availability.

**Figure 4:**
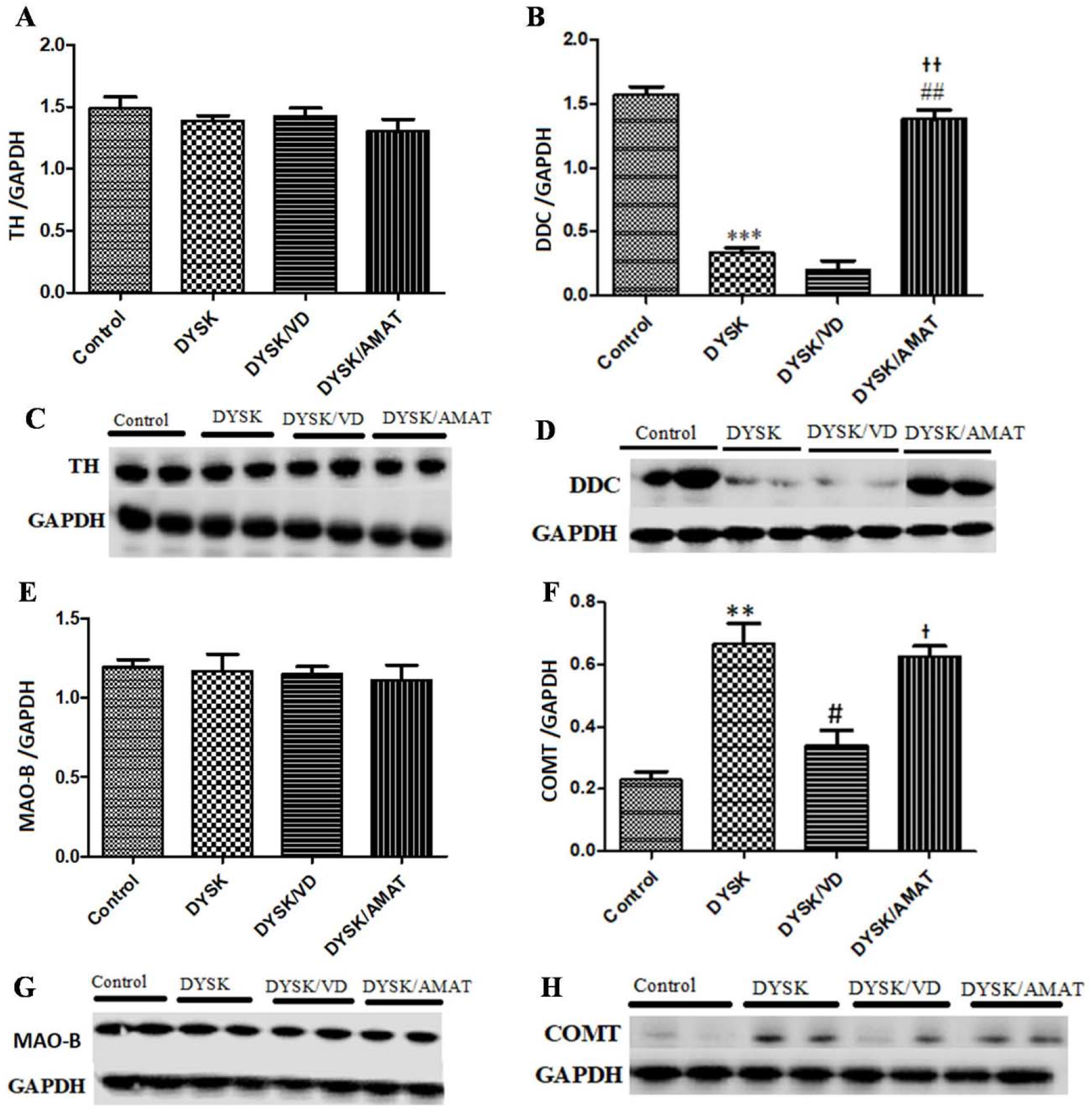
VD_3_ did not affect the expression of proteins involved in dopamine metabolism following chronic administration of L-DOPA in PD mice. Intervention with VD_3_ had no effect on the expression of TH (A), DDC (B) and MAO-B (E); reduced the expression of COMT (F). Quantification of band density of blots and the representative blots showing the expression of TH (C), DDC (D), MAO-B (G) and COMT (H) in the striatum. GAPDH was used as internal control. TH; tyrosine hydroxylase, DDC; dopamine decarboxylase, MAO-B; monoamine Oxidase-B, COMT; cathecol-O-methyl transferase. DYSK/VD; dyskinetic mice that received VD_3_, DYSK/AMAT; dyskinetic mice treated with amantadine. ^**^p<0.01, ^***^p<0.001 *vs* control; ^#^p<0.05, ^##^p<0.01 vs DYSK, ^+^p<0.05, ^++^p<0.01 vs DYSK/VD

### VD_3_ did not affect metabolic enzymes involved in dopamine neurotransmission in dyskinetic mice

As shown in Figure 4A, there was no significant difference in the expression of TH across all groups. As such, treatment of dyskinetic mice with VD_3_ did not alter the striatal level of TH [F (3, 7) = 0.99, p=0.48]. The expression of DDC was not changed following treatment with VD_3_ compared with dyskinetic mice whereas, amantadine significantly increased DDC expression compared with dyskinesia [Figure 4B: F (3, 7) = 128.5, p=0.0002] thereby promoting the release of dopamine. MAO-B and COMT catalyze the breakdown of dopamine. There was no significant difference in the expression of MAO-B across the groups, thus, neither intervention with VD_3_ nor amantadine altered the expression of MAO-B in dyskinetic mice (Figure 4E: F (3, 7) = 0.21, p=0.89). COMT was significantly increased in dyskinesia group compared with control. Treatment with amantadine had no significant effect on its expression while VD_3_ caused a significant decline in expression of COMT compared with dyskinesia group (Figure 4F: F (3, 7) = 21.71, p=0.0006).

### Anti-dyskinetic action of VD_3_ is linked to its modulatory effects on oxidative stress, microglial activation, inflammatory response, and apoptotic signaling

Oxidative stress, neuro-inflammation, microglial activation and apoptotic signaling are thought to be compromised in dyskinesia. Hence, we determined the impact of dyskinesia on these processes and the subsequent effect of VD_3_ intervention.

In Figure 5A, chronic L-DOPA-induced AIMs in dyskinetic mice was accompanied by the generation of ROS as shown by the marked expression of p47phox, (the main regulator and marker of NADPH oxidase which consequently drives ROS production). Interventions with VD_3_ (P<0.01) or amantadine (p<0.05) significantly attenuated the expression of p47phox compared with dyskinesia group, thus, a reduction in oxidative stress. There was also a significant upregulation of p47phox in amantadine group compared with VD_3_ mice showing that administration of VD_3_ was able to reduce production of ROS than amantadine in dyskinesia. Figure 5A: F (3, 7) = 77.50, p=0.0005.

**Figure 5:**
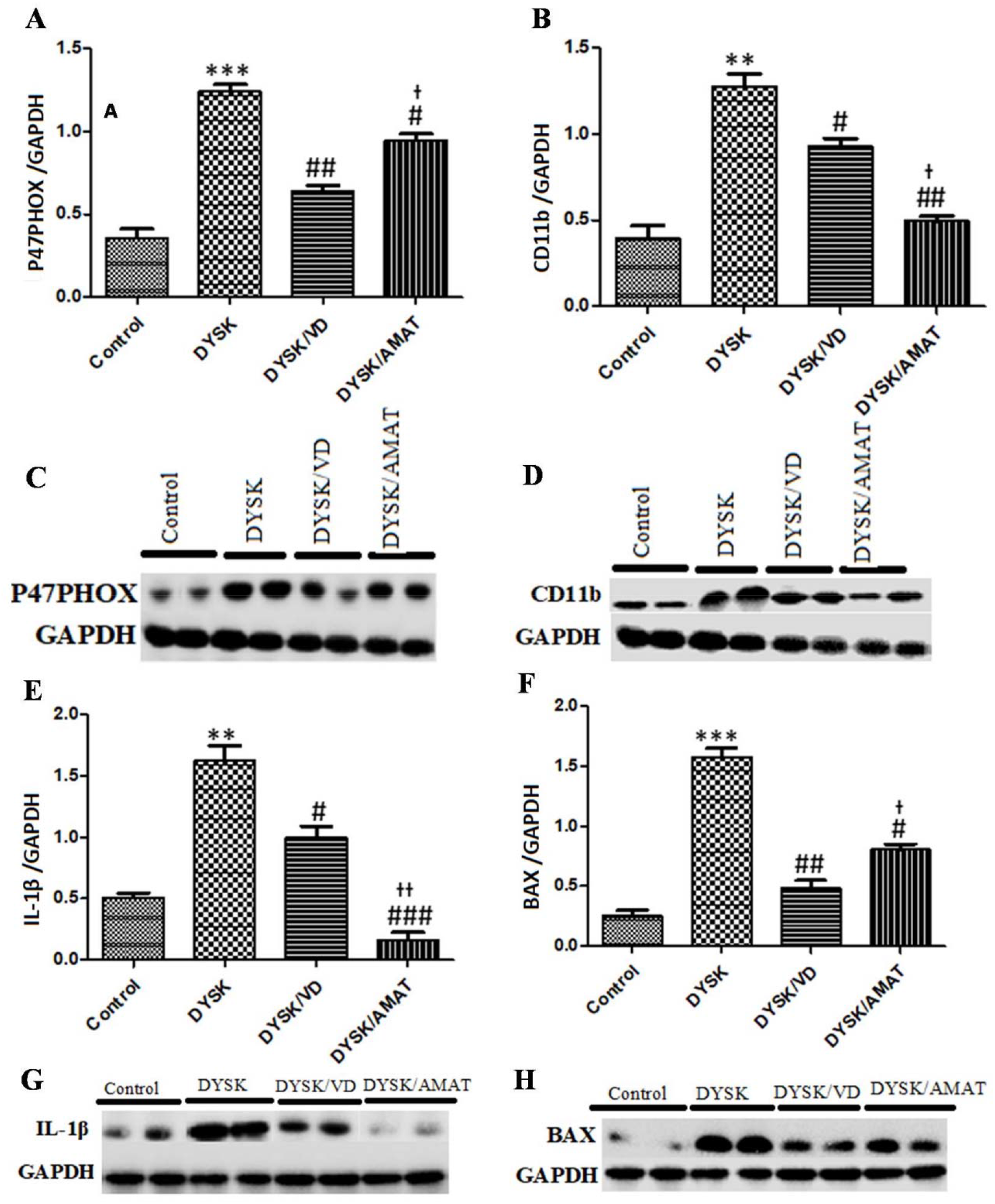
VD_3_ altered the expression of markers of oxidative stress, microglial, inflammation and apoptosis following chronic administration of L-DOPA in PD mice. Administration of VD_3_decreased the expression of p47phox (A), CD11b (B), IL-1β (E) and BAX (F). Quantification of band density of blots and the representative blots showing the expression of p47phox (C), CD11b (D), IL-1β (G) and BAX (H) in the striatum. GAPDH was used as internal control. p47phox; phagocyte NADPH oxidase organizer, CD11b; marker of microglial, IL-1β; Interleukin 1β, BAX; Bcl-2-associated X protein. DYSK/VD; dyskinetic mice that received VD_3_, DYSK/AMAT; dyskinetic mice treated with amantadine. ^**^p<0.01, ^***^p<0.001 *vs* control; ^#^p<0.05, ^##^p<0.01, ^###^p<0.001 *vs* DYSK, ^+^p<0.05, ^++^p<0.01 vs DYSK/VD

CD11b, a surface marker expressed by activated microglial was markedly expressed in dyskinetic mice compared to control mice. Intervention with either VD_3_ or amantadine significantly attenuated CD11b. Amantadine was better at reducing microglial activation than VD_3_ as shown by a significant decrease in the level of CD11b in the amantadine group compared with VD_3_ group. [Figure 5B: F (3, 7) = 50.58, p=0.0012].

Prolonged treatment with L-DOPA has been reported to increase the risk of inflammatory activity both in patients and animal models of PD. ^7^ In Figure 5E, chronic administration of L-DOPA led to increased expression of IL-Iβ, an inflammatory cytokine, in dyskinetic group of mice when compared with control. Treatment with VD_3_ for 14 days attenuated the expression of IL-1β. Similarly, amantadine significantly reduced IL-1β compared with dyskinesia and VD_3_ mice respectively [Figure 5E: F (3, 7) = 57.49, p=0.001].

Based on Figure 5F, *Striatal* expression of BAX, a pro-apoptotic member of Bcl-2 family and an important regulator of apoptosis was significantly upregulated in dyskinesia compared with control. Intervention with VD_3_ attenuated BAX expression. Administration of amantadine also reduced BAX compared with dyskinesia group but its effect was lower when compared with VD_3._ Hence, the anti-apoptotic activity of VD_3_ was more pronounced than amantadine. [Figure 5F: F (3, 7) = 95.11, p=0.0004].

## Discussion

The hemi-parkinsonian mouse model of LID is a valuable and unique tool in dyskinesia research, because it allows interspecies comparison of drug response, is suitable for genetic manipulation and is particularly advantageous in target validation. Unilaterally lesioned mice that were chronically treated with L-DOPA (20 mg/kg plus 12.5mg/kg of benserazide) developed a stable degree of dyskinesia by the sixth day of treatment, scoring the maximal cumulative ALO AIMs value on day 8. ALO AIMs developed gradually and progressively having a similar temporal pattern, with maximal dyskinetic expression after 6 days of chronic treatment with L-DOPA. This may reflect the homogeneity of the lesion within the dorsolateral striatum as this region is documented to receive somatotopic cortical projections representing the trunk, forepaw and orofacial muscles. ^47^ Our result is slightly different from the report of Bido *et al*., wherein chronic administration of L-DOPA to Parkinsonian mice showed maximal expression of dyskinetic behavior after 5 days of treatment. Similar to their report, we noted the appearance of AIMs as early as 20 mins post L-DOPA treatment. In contrast, maximal dyskinetic response was observed at 60 mins, followed by a decline afterwards, compared with a maximal expression observed by Bido *et al*., at 40 min after which expression declined. ^48^ The reason for the slight disparity in the two results may be attributable to the difference in the dyskinetic dose of L-DOPA treatment. Bido *et al*., administered 15 mg/kg for inducing dyskinesia while the present study used 20 mg/kg, although a duration of 10 days of L-DOPA treatment was used in both studies. Genetic interference may also contribute to the difference in results, since the two studies used different strains of mice. A dose of 40 mg/kg of amantadine (reference drug) was used because it has been proven to be effective in reducing ALO AIMs in rodents without affecting the locomotive component of AIMs, ^39, 45^ which is also considered a marker of the therapeutic effect of L-DOPA. ^46^ In addition, orolingual AIM subtype had the least score while axial appearance was the most represented with the highest score, which is in line with prior studies. ^14, 20, 48^ VD_3_ showed anti-dyskinetic effects as it produced a significant attenuation of global AIMs on days 11 and 14, which corresponds to 22% and 32.7% reduction in dyskinetic movements respectively. However, the reference drug amantadine markedly attenuated ALO AIMs progressively throughout the observation period with the highest percentage ALO AIMs reduction of 64.5%. Therefore, VD_3_ attenuated L-DOPA-induced AIMs, but its anti-dyskinetic action was less than amantadine. There is no report yet on the role of VD_3_ on LID, whereas other studies have reported the anti-dyskinetic effect of VD and vitamin E in reducing the threshold of Tardive dyskinesia, ^33, 34^ a side effect of prolonged use of anti-psychotic drugs. Dyskinesia refers to any involuntary movement, such as chorea, tic, dystonia, ballism that affect any part of the body. A link reportedly occurs between occurrence of essential tremor and vitamin D receptor polymorphism. As such, the rs2228570 variant of the vitamin D receptor gene has been associated with sporadic essential tremor. ^49^

Vitamin D has been associated with many distinct neurological functions, particularly neuroprotection. Insufficient levels have consequently been linked to an increased likelihood of developing neurological diseases such as movement disorders. On the contrary, a clinical study which investigated the effect of VD supplementation in dyskinetic patients reported that VD_3_ did not improve dyskinesia rating when compared to placebo. ^50^

Many pathways such as serotonergic, cholinergic and glutamatergic pathways were reportedly involved in the uncontrolled release of dopamine and the abnormal involuntary movements in dyskinesia.^1, 51^ The involvement of *striatal* DA synthetic (TH and DDC) and catabolic (MAO-B and COMT) enzymes was assessed and the effect of VD_3_ on these regulatory enzymes was investigated in dyskinetic mice. TH, the rate limiting enzyme in the synthesis of dopamine, was unaffected in dyskinetic mice compared with control group. This is in contrast but has similar implication with the report of Pons *et al*. ^52^ which showed a deficiency in TH in LID. The interpretation of these two reports is that either a decrease or no effect of TH in dyskinetic mice does not support the accompanied rise in DA level. This is because TH converts tyrosine to L-DOPA and as such any event that will promote DA synthesis should positively enhance the activity of TH. Similarly, the expression of MAO-B in the dyskinetic mice was not different from those of the control mice. Hence, based on the data from this study, both enzymes are not altered in DA transmission in dyskinetic mice. As such, the reported high level of DA that accompanies dyskinesia is not likely to be from this source.

The present study shows that DDC expression was significantly reduced in the dyskinesia group compared to the control mice. The observed decrease in DDC indicates that the reported high level of *striatal* DA that accompanies dyskinesia ^53^ could not have resulted from the activity of presynaptic *striatal* DDC. The expression of DDC in this study does not support DA release. Thus, these findings corroborate the assertion that the typical high DA level in dyskinesia could have resulted from extra-dopaminergic sources. ^54^ The activity of COMT was significantly increased in dyskinesia mice compared with the control group. If the level of DA in dyskinesia were to be from the conventional pathway of DA metabolism, then COMT level should have been reduced or unaltered. That is, DA break down ought to be inhibited thereby enhancing DA availability. In contrast, the reverse was observed which did not support DA availability. Reports have also shown that COMT could be an independent genetic predisposition factor for the development and severity of LID. Solís *et al*.,^55^ showed that overexpression of COMT in transgenic mice increases susceptibility to LID and potentiates expressions of molecular correlates of dyskinesia (FosB and pAcH3 expression). The regulatory role of COMT in the development and severity of LID is supported by the fact that COMT inhibitors have been employed clinically to reduce LID through enhanced L-DOPA bioavailability and prolonged DA action in the *striatum*. ^56^ Genetic variations in the COMT genes is being investigated and suggested as the underlying factor for the large disparity in the susceptibility of PD patients to LID and response to treatment. ^57, 58^

Treatment of dyskinetic mice with either VD_3_ or Amantadine did not alter the *striatal* level of TH. Similarly, the expression of DDC was not significantly altered following treatment with VD_3_ compared with dyskinetic mice whereas, amantadine significantly increased DDC expression, thereby promoting the release of dopamine. One of the mechanisms through which amantadine acts is by inhibiting DA reuptake ^59, 60^ and increasing DDC activity ^61, 62^ which is in line with the findings of this study. However, this effect of amantadine is difficult to reconcile with its clinical use as an anti-dyskinetic drug because a potentiation of the L-DOPA-induced DA release ^63^ would also lead to stimulation of D1 receptors on the *striatal* cell bodies and *nigral* terminals of *striato-nigral* GABA neurons, thereby promoting LID. To our knowledge, this is the first report on the effect of VD_3_ on dopamine metabolic enzymes in L-DOPA-induced dyskinesia. There was no significant difference in the expression of MAO-B across the groups, neither did intervention with VD_3_ nor amantadine alter the expression of MAO-B in dyskinetic mice. In contrast, treatment with VD_3_ dampened the expression of COMT while amantadine had no significant effect on its expression. While a recent report attributed the neuroprotective impact of VD_3_ in mice model of PD to its ability to modulate proteins involved in dopamine metabolism, ^38^ on the contrary, it appears that the mechanism of anti-dyskinetic effect of VD_3_ (as shown in the present study) does not involve the enzymatic regulation of dopamine metabolism. More studies are needed to further unravel the involvement of VD_3_ in dyskinesia.

The pathophysiological process by which vitamin D deficiency might lead to the development and progression of movement disorders, is not yet fully understood. High concentrations of vitamin D metabolites and of vitamin D receptor proteins are found in the basal ganglia and connected structures. ^64^ In the basal ganglia, vitamin D has been shown to function as a modulator in brain development and as a neuroprotectant. ^64^ Another important mechanism is that vitamin D has exhibited an association with the regulation of the synthesis of nerve growth factors that is responsible for the growth and survival of neurons. ^65^

Chronic LIDs in mice may be accompanied by the generation of ROS as indicated by the marked expression of p47phox, a marker and organizer of NADPH oxidase, which in turns drives ROS via super oxide production). ^66, 67^ Interventions with VD_3_ significantly attenuated the expression of p47phox which depicts a reduction in the activities of ROS, thereby inhibiting oxidative stress. Vitamin D_3_ reportedly inhibited the synthesis of inducible nitric oxide synthase, an enzyme induced in neurons and non-neuronal cell during ischemia and in neurodegenerative conditions, which catalyzes nitric oxide, a free radical that can damage cells. It stimulates γ-glutamyl transpeptidase activity which is important in the synthesis of glutathione, antioxidant that neutralizes free radicals and in this way protects cells from damage. ^68, 69^ Moreover, the inhibitory action of VD_3_ on ROS and microgial activation is well posited in literatures. ^70-72^ Vitamin D has been reported to act through several other mechanisms, including effects on protein expression, oxidative stress, inflammation, and cellular metabolism. ^73, 74^

CD11b is the most important and potent surface marker expressed by activated microglia. ^75^ It was markedly expressed in dyskinetic mice compared to the control mice showing that microglia activation is involved in the progression of dyskinesia. Intervention with either VD_3_ or amantadine significantly attenuated CD11b. Prolonged treatment with L-DOPA has been reported to increase the risk of inflammatory activities both in patients and animal models of PD. ^7^ Chronic administration of L-DOPA led to increase in the expression of IL-Iβ, an inflammatory cytokine in dyskinetic group of mice compared with control. Treatment with VD_3_ for 14 days attenuated the level of IL-1β. Similarly, amantadine reduced inflammation significantly compared with dyskinesia and VD_3_ mice respectively. Intervention with VD_3_ markedly attenuated IL-1β-induced inflammatory activity) and apoptotic signaling. VD reportedly upregulated Bcl-2, an apoptosis inhibitor in a non-pathologic condition and exerted anti-apoptotic effect in ovarian cancer cells by inhibiting apoptosis mediated by death receptors. ^76^

Striatal expression of BAX, a pro-apoptotic member of Bcl-2 family and an important regulator of apoptosis was significantly upregulated in dyskinesia compared with control. Intervention with VD_3_ attenuated BAX level. Administration of amantadine also reduced apoptotic signaling compared with the dyskinesia group but its effect was lower when compared with VD_3_. Hence, the anti-apoptotic activity of VD_3_ was more pronounced than amantadine. Hypovitaminosis D has been shown to cause apoptosis by diminishing the expression of cytochrome C, thereby decreasing the cell cycle of neurons. ^69^

## Conclusion

Vitamin D_3_ showed anti-dyskinetic effects by attenuating dyskinetic AIMs. It did not alter the expression of the major enzymes involved in dopaminergic neurotransmission, hence, its anti-dyskinetic effect does not involve dopaminergic modulation. The anti-dyskinetic action of VD_3_ could have resulted from its modulatory effect on the generation of reactive oxygen species, microglial responses, inflammation, and apoptotic signaling. Therefore, its therapeutic use in the management of dyskinesia is suggested. More studies are required to further evaluate these findings.

## Acknowledgement

The authors would like to appreciate Mr Ishan Methrota who assisted with part of the laboratory work.

## Authors’ Contributions

OMO, AA and BVO designed the research, BAA performed the experiment and analyzed the data, BAA, AA and ABN wrote the manuscript, BVO and CCL read, edited and approved the manuscript.

